# Frontoparietal network dynamics impairments in juvenile myoclonic epilepsy revealed by MEG energy landscape

**DOI:** 10.1101/703074

**Authors:** Dominik Krzemiński, Naoki Masuda, Khalid Hamandi, Krish D Singh, Bethany Routley, Jiaxiang Zhang

**Author notes:** Correspondence should be addressed to: Dominik Krzemiński or Jiaxiang Zhang.

## Abstract

Juvenile myoclonic epilepsy (JME) is a form of idiopathic generalized epilepsy affecting brain activity. It is unclear to what extent JME leads to abnormal network dynamics across functional networks. Here, we proposed a method to characterise network dynamics in MEG resting-state data, combining a pairwise maximum entropy model (pMEM) and the associated energy landscape analysis. Fifty-two JME patients and healthy controls underwent a resting-state MEG recording session. We fitted the pMEM to the oscillatory power envelopes in theta (4-7 Hz), alpha (8-13 Hz), beta (15-25 Hz) and gamma (30-60 Hz) bands in three source-localised resting-state networks: the frontoparietal network (FPN), the default mode network (DMN), and the sensorimotor network (SMN). The pMEM provided an accurate fit to the MEG oscillatory activity in both patient and control groups, and allowed estimation of the occurrence probability of each network state, with its regional activity and pairwise regional co-activation constrained by empirical data. We used energy values derived from the pMEM to depict an energy landscape of each network, with a higher energy state corresponding to a lower occurrence probability. When comparing the energy landscapes between groups, JME patients showed fewer local energy minima than controls and had elevated energy values for the FPN within the theta, beta and gamma-bands. Furthermore, numerical simulation of the fitted pMEM showed that the proportion of time the FPN was occupied within the basins of characteristic energy minima was shortened in JME patients. These network alterations were confirmed by a significant leave-one-out classification of individual participants based on a support vector machine employing the energy values of pMEM as features. Our findings suggested that JME patients had altered multi-stability in selective functional networks and frequency bands in the frontoparietal cortices.

**Highlights:**

- An energy landscape analysis characterises the dynamics of MEG oscillatory activity
- Patients with JME exhibit fewer local minima of the energy in their energy landscapes
- JME affects the network dynamics in the frontoparietal network.
- Energy landscape measures allow good single-patient classification.

## 1 Introduction

Juvenile myoclonic epilepsy (JME) is the most common syndrome of the wider group of idiopathic generalized epilepsies (Wolf and Beniczky, 2014). Patients with JME often exhibit three main types of seizures: myoclonic, absence and generalized tonic-clonic seizures (Wolf et al., 2015a). Typical JME characteristics are normal or close to normal clinical MRI of the brain and interictal EEG with irregular spike-waves or polyspike-waves with frontal predominance (Camfield et al., 2013). JME patients are susceptible to seizure precipitation after sleep deprivation, alcohol usage, excise or demanding cognitive processing (Delgado-Escueta and Enrile-Bacsal, 1984; Yacubian and Wolf, 2014). JME is a lifelong condition and treatment with antiepileptic drugs is usually necessary.

Although the pathogenetic mechanisms of JME is still not fully understood (Berkovic et al., 1998), JME has been recognised as a network disorder affecting brain activity and connectivity that leads to cognitive impairments (Chowdhury et al., 2014; Wolf et al., 2015a) and personality traits similar to patients with frontal lobe lesions (Engel J, 1997). BOLD functional MRI (fMRI) and diffusion weighted imaging showed hyper-connectivity in the frontal lobe in JME (Vollmar et al., 2012; Caeyenberghs et al., 2015). Electrophysiological data suggests that JME has an impact on multiple functional networks, including the frontoparietal network (FPN) (Wolf et al., 2015a), the default mode network (DMN) (McGill et al., 2012), and the sensorimotor network (SMN) (Clemens et al., 2013), which may be driven by dysfunctional thalamocortical circuitry (Gotman et al., 2005; Hamandi et al., 2006; Betting et al., 2006; Kim et al., 2007a).

Several sensitive markers from resting EEG and magnetoencephalography (MEG) recordings have been identified for classifying patients with epilepsy and predicting seizure onsets, including information entropy (Kannathal et al., 2005; Song et al., 2012; Song and Zhang, 2013), Lyapunov exponent (Iasemidis et al., 1990; Babloyantz and Destexhe, 1986) and phase plane portraits (Iasemidis et al., 1990). These methods describe statistical regularities of electrophysiological signals from a dynamical system perspective, in line with the theoretical account of epileptic seizures as bifurcations from stable states (da Silva et al., 2003). In JME, however, it is yet unclear to what extent JME patients had atypical network dynamics across different functional networks and whether the abnormalities of network dynamics are frequency specific.

This study addressed these questions by applying a novel pairwise maximum entropy model (pMEM) approach (Yeh et al., 2010) to source-localised, frequency-specific MEG resting-sate oscillatory activity. The pMEM is a statistical model of the occurrence probability of network states, with its parameters being constrained by the network’s regional activity and pairwise regional co-activation from empirical data. According to the principle of maximum entropy, the pMEM is the most parsimonious second-order model of a system with minimum assumptions (Jaynes, 1957). The pMEM has been successfully applied to the collective behaviour of spiking neural networks (Tkacik et al., 2006; Schneidman et al., 2006; Tang et al., 2008) and BOLD fMRI responses (Watanabe et al., 2013, 2014; Ashourvan et al., 2017; Ezaki et al., 2018). Here, we extended this theoretical framework to MEG oscillatory activity in three functional networks: FPN, DMN and SMN. Furthermore, based on the fitted pMEM to individual participants, we depicted an energy landscape for each of the networks at theta (4-7 Hz), alpha (8-13 Hz), beta (15-25 Hz) and gamma (30-60 Hz) bands. The energy landscape is a graphical representation of all network states and their energy values (Ezaki et al., 2017). We then compared several quantitative measures obtained from the energy landscapes between JME patients and controls.

Our results demonstrated that the pMEM provided a good fit to the statistical properties of functional networks in both JME and control groups. JME patients showed reduced numbers of local energy minima and elevated energy values in the theta, beta and gamma-band FPN activity, but not in the SMN. We further demonstrated that the pMEM could be used as a generative model for simulating dysfunctional network dynamics in JME, and as a predictive model for single-patient classification. These findings suggest anatomically- and frequency-specific network abnormalities in JME.

## 2 Methods

### 2.1 Participants

Fifty-two subjects participated in the experiment. Demographic and clinical features of the participants are summarized in Table 1. Twenty-six patients with JME were recruited from a specialist clinic for epilepsy at University Hospital of Wales in Cardiff. Consensus clinical diagnostic criteria for JME were used by an experienced neurologist (Trenité et al., 2013). Inclusion criteria were: (1) seizure onset in late childhood or adolescence with myoclonic jerks, with or without absence seizures, (2) generalised tonic-clonic seizures, (3) normal childhood development as assessed on clinical history and (4) generalised spike wave on EEG and normal structural MRI. Twenty-six healthy control participants with no history of significant neurological or psychiatric disorders were recruited from the regional volunteer panel. All testing was performed with participants’ taking their usual medication. The study was approved by the South East Wales NHS ethics committee, Cardiff and Vale Research and Development committees, and Cardiff University School of Psychology Research Ethics Committee. Written informed consent was obtained from all participants.

**Table 1:** Demographics of patients with JME and and healthy control participants. (MJ - myoclonic jerks, GTCS - generalised tonic clonic seizures, LEV - leveiracetam, VPA - sodium valproate, LTG - lamotrigine, TPM - topiramate, ZNM - zonisamide.)

|  | **Patients** | **Controls** |
| --- | --- | --- |
| Number of participants | 26 (8 males) | 26 (7 males) |
| Age median | 27 | 27 |
| Age range | 19 - 45 | 18 - 48 |
| Seizure type<br>(number of patients) | MJ (26)<br>Absences (15)<br>GTCS (26) | - |
| Anti-epileptic drugs<br>(Number of patients taking the drug) | LEV (13), VPA (12),<br>LTG (5), TPM (4),<br>ZNM (4) | - |

### 2.2 MEG and MRI data acquisition

All participants underwent separate MEG and MRI sessions. Whole-head MEG recordings were made using a 275-channel CTF radial gradiometer system (CTF Systems, Canada) at a sampling rate of 600 Hz. An additional 29 reference channels were recorded for noise cancellation purposes and the primary sensors were analysed as synthetic third-order gradiometers (Vrba and Robinson, 2001). Up to three sensors were turned off during recording due to excessive sensor noise. Subjects were instructed to sit comfortably in the MEG chair while their head was supported with a chin rest and with eyes open focus on a red dot on a grey background. For MEG/MRI co-registration, fiduciary markers that are identifiable on the subject’s anatomical MRI were placed at fixed distances from three anatomical landmarks (nasion, left and right preauricular) prior to the MEG recording, and their locations were further verified using high-resolution digital photographs. The locations of the fiduciary markers were monitored before and after MEG recording. Each recording session lasted approximately 5 minutes.

Whole-brain T1-weighted MRI data were acquired using a General Electric HDx 3T MRI scanner and a 8-channel receiver head coil (GE Healthcare, Waukesha, WI) at the Cardiff University Brain Research Imaging Centre with an axial 3D fast spoiled gradient recalled sequence (echo time 3 ms; repetition time 8 ms; inversion time 450 ms; flip angle 20°; acquisition matrix 256×192×172; voxel size 1×1×1 mm).

### 2.3 Data pre-processing

Continuous MEG data was first segmented into 2 s epochs. Every epoch was visually inspected. Epochs containing major motion, muscle or eye-blink artefact, or interictal spike wave discharges were excluded from subsequent analysis. The artefact-free epochs were then re-concatenated. This artefact rejection procedure resulted in cleaned MEG data with variable lengths between 204 s and 300 s across participants, and the data lengths were comparable between JME patients and controls (*t*(50) = 1.38, *p* = 0.17). The 200 s of cleaned MEG data was used in subsequent analysis. For participants with longer than 200 s cleaned MEG data, a continuous segment of 200 s during the middle of recording session was used.

### 2.4 Source localization of oscillatory activity in resting-state networks

We analysed the MEG oscillatory activity using an established source localisation method for resting-state networks (Brookes et al., 2011; Hall et al., 2013; Muthukumaraswamy et al., 2013). For each participant, the structural MRI scan was co-registered to MEG sensor space using the locations of the fiducial coils and the CTF software (MRIViewer and MRIConverter). The structural MRI scan was segmented and a volume conduction model was computed using the semi-realistic model (Nolte, 2003). The preprocessed MEG data was band-passed filtered into four frequency bands: theta 4-8 Hz, alpha 8-12 Hz, beta 13-30 Hz, and low-gamma 35-60 Hz (Niedermeyer, 2005). For each frequency band, we downsampled the data to 250 Hz and computed the inverse source reconstruction using an LCMV beamformer on a 6-mm template with a local spheres forward model in Fieldtrip (version 20161101, http://www.fieldtriptoolbox.org). The atlas-based source reconstruction was used to derive virtual sensors for every voxel in each of the 90 regions of the Automated Anatomical Label (AAL) atlas (Hipp et al., 2012). Each virtual sensor’s time course was then reconstructed.

We focused our analysis on three resting-state networks (Fig. 2): the fronto-parietal network (FPN), the default mode network (DMN), and the sensorimotor network (SMN) in which electrophysiological changes had been reported in patients with epilepsy (McGill et al., 2012; Clemens et al., 2013; Wolf et al., 2015a). Each resting-state network comprised of bilateral regions of interest (ROIs) from the AAL atlas identified in previous studies (Tewarie et al., 2013; Rosazza and Minati, 2011). The FPN included middle frontal gyrus (MFG), pars triangularis (PTr), inferior parietal gyrus (IPG), superior parietal gyrus (SPG) and angular gyrus (AG). The DMN included orbitofrontal cortex (OFC), anterior cingulate cortex (ACC), posterior cingulate cortex (PCC), precuneus (pCUN) and AG. The SMN included precentral gyrus (preCG), postcentral gyrus (postCG), supplementary motor area (SMA). For each ROI, its representative time course was obtained from the voxel in that ROI with the highest temporal standard-deviation. The mean MEG activities of the ROIs of each network were not significantly different between JME patients and controls (FPN: *F*(1, 50) = 0.75, *p* = 0.39; DMN: *F*(1, 50) = 0.21, *p* = 0.65; SMN: *F*(1, 50) = 0.15, *p* = 0.70).

To calculate the oscillatory activity, we applied Hilbert transformation to each ROI’s time course, and computed the absolute value of the analytical representation of the signal to generate an amplitude envelope of the oscillatory signals in each frequency band.

### 2.5 Pairwise maximum entropy model of MEG oscillatory activity

During rest, different brain regions exhibit pairwise co-occurrence of oscillatory activity (Horwitz, 2003) and rapid changes of brain network states (Stam and Straaten, 2012). To obtain an estimate of network state transitions and their probabilities, we fitted a pMEM to individual participant’s MEG data, separately for each resting-state network and each frequency band.

According to the principle of maximum entropy, among all probabilistic models describing empirical data, one should choose the one with the largest uncertainty (i.e., entropy), because it makes the minimum assumptions of additional information that would otherwise lower the uncertainty (Yeh et al., 2010). The pMEM estimates the state probability of a network, with its regional activity and regional co-occurrence to be constrained by empirical data. Previous studies have successfully used the pMEM to accurately measure network dynamics in neural recording (Tkacik et al., 2006) and functional MRI data (Watanabe et al., 2013). The current study used a similar approach for modelling MEG oscillatory activity (Fig. 1). Below we outlined the theoretical background and the fitting procedure. A more detailed description of the pMEM modelling and subsequent energy landscape analysis is available elsewhere (Ezaki et al., 2017).

**Figure 1:**
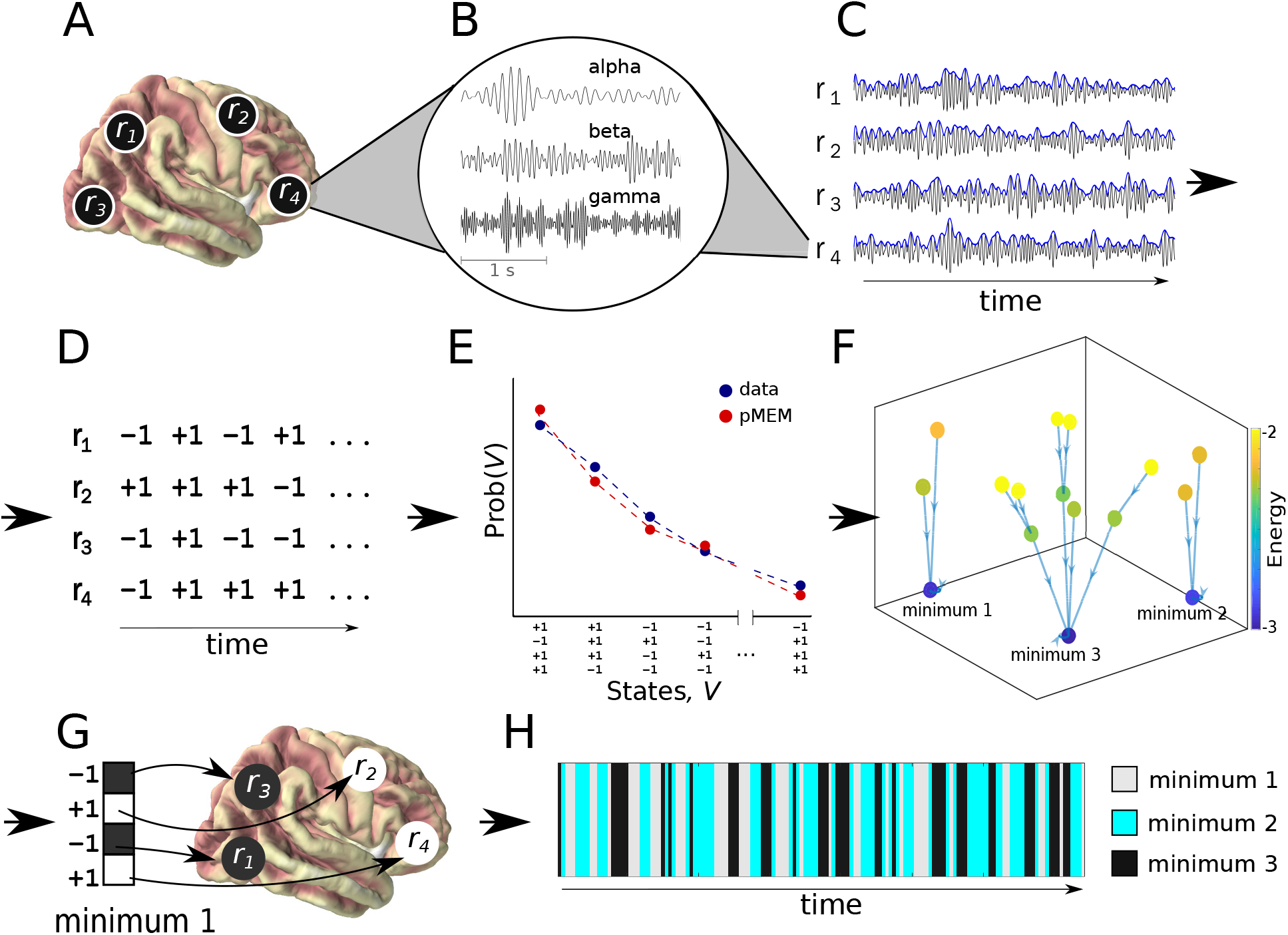
Illustration of the energy landscape analysis on a network of 4 ROIs. **A**. Selection of ROIs from the source-space signals. **B**. Signal filtering in frequency bands of interest. **C**. Envelope extraction using the absolute value of the analytical representation of the signal. **D**. Binarisation of the data; **E**. Fitting the pMEM to match the empirical data distribution of binarised netwrok states. **F**. Determining the relationships between network states using the Dijkstra algorithm on energy values. **G**. Interpretation of local minima of the energy on the anatomical level. **H**. Simulation of the occurrence of network states belonging to different basins.

Consider a resting-state network consisting of *N* ROIs. For each real-valued ROI’s signal, we thresholded the ROI’s Hilbert envelope according to the median of the amplitude. Data points above the threshold were denoted as high oscillatory power (+1), and data points below the threshold were denoted as low oscillatory power (−1). The oscillatory activity in ROI *i* (*i* = 1,…, *N*) at time *t* was transformed to a binary time series *r_i_*(*t*), with *r_i_*(*t*) = +1 for high oscillatory activity and *r_i_*(*t*) = −1 for low oscillatory activity. The activity pattern of a *N*-dimensional binary vector **s**(*t*) = [*r*_1_(*t*), *r*_2_(*t*), …, *r_N_*(*t*)], representing the state of the network at time *t*.

The *N*-ROI network has a total of 2^*N*^ possible states **s**_*k*_ (*k* = 1,…, 2^*N*^). From the binarized oscillatory activity, we calculated the probability of occurrence of each network state, denoted by *P*_emp_(**s**_*k*_). We further calculated the empirical average activation rate for each ROI 〈*r_i_*〉_emp_ and the pairwise co-occurrence between any two ROIs 〈*r_i_r_j_*〉_emp_:

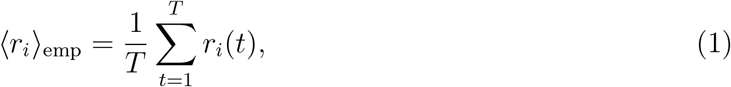

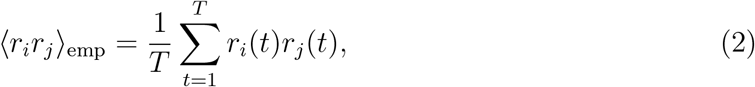

where *T* denotes the number of timepoints in the data. The fitting procedure aimed to identify a pMEM model that preserves the constraints in Equations (1) and (2) and reproduces the empirical state probability *P*_emp_(**s**_*k*_) with the maximum entropy. It is known that the pMEM follows the Boltzman distribution (Yeh et al., 2010), given by

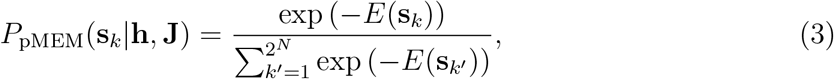

where *E*(**s**_*k*_) represents the energy of the network state **s**_*k*_, defined by

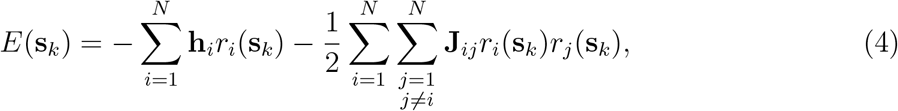

and *r_i_*(**s**_*k*_) refers to the *i*-th element of the network state **s**_*k*_. **h** and **J** are the model parameters to be estimated from the data: **h** = [*h*_1_, *h*_2_, …, *h_N_*] represents the bias in the intensity of the oscillatory activity in each ROI; **J** = [*J*_11_, *J*_12_, …, *J_NN_*] represents the coupling strength between two ROIs. The average of the activation rate 〈*r_i_*〉_mod_ and pairwise co-occurrence 〈*r_i_r_j_*〉_mod_ expected by the pMEM are given by:

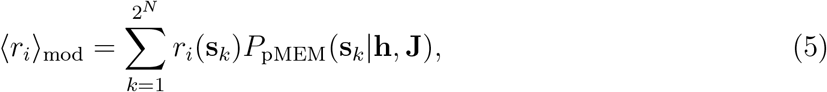

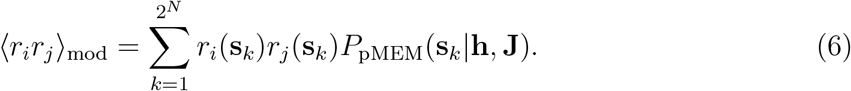

We used a gradient ascent algorithm to iteratively update **h** and **J**, until 〈*r_i_*〉 _mod_ and 〈*r_i_r_j_*〉_mod_ match 〈*r_i_*〉_emp_ and 〈*r_i_r_j_*〉_emp_ from the observed data, with a stop criterion of 5 × 10^6^ steps. In each iteration, the updates of the parameters were given by 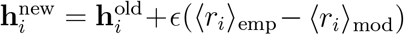 and 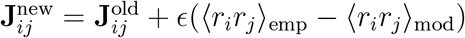. The learning rate *ϵ* was set to 10^−8^.

As in previous studies (Watanabe et al., 2014; Ezaki et al., 2017, 2018), we used an accuracy index:

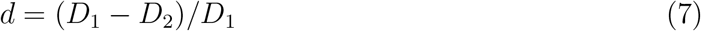

to quantify the goodness of fit of the pMEM, where

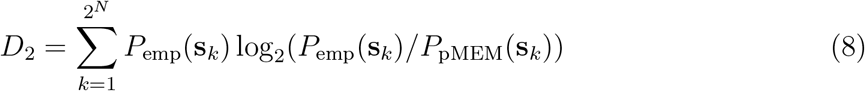

is the Kullback-Leibler divergence between the probability distribution of the pMEM and the empirical distribution of the network state. *D*_1_ represents the Kullback-Leibler divergence between the independent MEM and data. By definition, the independent MEM is restricted to have no pairwise interaction (i.e., **J** = 0). Therefore, *d* represents the surplus of the fit of the pMEM over the fit of the independent model. The index *d* =1 when the pMEM reproduces the empirical distribution of activity patterns and interactions without errors, and *d* = 0 when the pairwise interactions do not contribute to the description of the empirical distribution.

### 2.6 Energy landscape of resting-state network dynamics

The pMEM parameters **h** and **J** determine the energy *E*(**s**_*k*_) of each network state **s**_*k*_ (*k* = 1,…, 2^*N*^), given by Equation (4). It is worth noting that, the current study used pMEM as a statistical model to be constructed from the MEG data, not as its literal notion from statistical physics. We did not claim that *E*(**s**_*k*_) represents the metabolic or physical energy of a biological system. Instead, the concept of the energy of a resting-state network stems from the information theory (Ezaki et al., 2017). Here, *E*(**s**_*k*_) indicates the model prediction of the inverse appearance probability of the state **s**_*k*_ under the empirical constraints of regional activity (parameter **h**) and regional interactions (parameter **J**). For instance, if *E*(**s**_*i*_) < *E*(**s**_*j*_), the pMEM predicts that the network activity pattern is more likely to be at the state **s**_*i*_ than **s**_*j*_.

For each resting-state network and each frequency band, we depicted an energy landscape as a graph of the energy function across the 2^*N*^ possible network states **s**_*k*_, characterising state probabilities and state transitions from the perspective of attractor dynamics (Watanabe et al., 2014). The energy landscape of a network was defined by two factors: the energy *E*(**s**_*k*_) of each network state, and an adjacency matrix defining the connectivity between network states. Two states were defined to be adjacent, or directly connected, if and only if just one ROI of the network had different binarized oscillatory activity (high vs. low). In other words, two states are adjacent when they have a Hamming distance of 1 between their binary activity vectors. For example, for a network with 4 ROIs, states [−1, −1, −1, +1] and [−1, −1, +1, +1] are adjacent, and states [−1, −1, −1, +1] and [−1, −1, +1, −1] are not.

### 2.7 Quantitative measures of energy landscape

We used three measures to understand the differences in the energy landscape between JME patients and healthy controls: (1) the number of local energy minima, (2) the relative energy of the local minima, and (3) the generative basin duration at significant minima.

#### 2.7.1 Number of local energy minima

A local energy minimum was defined as the network state with a lower energy value than all its adjacent neighbouring states. Because lower energy corresponds to a higher probability of occurrence, network states of local minima can be likened as attractors in attractor dynamics. For each participant, we exhaustively searched through the 2^*N*^ network states to identify all the local minima of the participant’s energy landscape. We then compared the number of local energy minima between JME and control groups (Fig. 4).

#### 2.7.2 Relative energy of the local minima

We calculated the mean energy 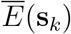 of each network state **s**_*k*_ averaged across all participants. Then, we used the mean energy to depict an aggregated landscape, which allowed us to identify common energy minima shared between JME patient and control groups. To test whether each local minimum in the aggregated energy landscape is a characteristic feature of the observed data, we conducted non-parametric permutation tests on the mean energy values. For each resting-state network and each frequency band, we calculated randomised mean energy values at each *s_k_* using 1000 permutations, with the pMEM parameters **h** and **J** of each of the participants unifomly randomly shuffled between ROIs and ROI pairs, respectively, in each permutation. This gave us a sampling distribution of the energy of each network state, under the null hypothesis that the energy values are not related to the observed oscillatory activities or observed pairwise regional co-occurrence. For each local minimum of the aggregated landscape, the level of significance (*p*-value) of that local minimum’s energy was estimated by the fraction of the permutation samples that were higher than the mean energy 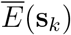 of that network state in the empirical data without shuffling. To account for the multiple statistical tests that were performed for all the local minima of each network, we evaluated the results using a Bonferroni-corrected threshold (*p*<0.05) for significance.

Because the shape of a energy landscape was partly determined by the global minimum (Watanabe et al., 2014; Ezaki et al., 2018), for each participant, we calculated the energy difference between a significant local minimum and the global energy minimum (i.e., the state with the lowest energy value on the landscape). We then compared this relative energy of the local minima between JME patients and healthy controls. From the networks with significant alternations of relative energy values in JME patients, we constructed a disconnectivity graph to describe clusters of local minima and the relationships between them (Becker and Karplus, 1997), where a cluster represents a group of local minima with high probabilities of subsequent occurrences (Ezaki et al., 2017).

#### 2.7.3 Basin duration at significant minima

On the aggregated energy landscape, the energy basin for each significant local minimum was identified using an existing method (Watanabe et al., 2014). We started at an arbitrary network state and moved downhill on the energy landscape to one of its neighbouring state with the lowest energy, until a local minimum was reached. The starting state is then assigned to the basin of the resulting local minimum. We repeated this procedure for all network states as the starting state.

We used the fitted pMEM as a generative model to simulate the dynamical changes in each resting-state network, and estimated the duration of the basin of each local minimum in the simulated dynamics. Similar to previous studies, we employed the Metropolis-Hastings algorithm to simulate time courses of network activity (Hastings, 1970). Each simulation started with a random network state **s**_*k*_. On each time step, one of the current state’s *N* neighbouring state **s**_*k*_ was selected with a probability of 1/*N* as the potential target of state transition, and the state transition occurred with a probability of exp [*E*(**s**_*k*_) − *E*(**s**_*k′*_)] when *E*(**s**_*k′*_) > *E*(**s**_*k*_) or 1 otherwise. For each participant, each network, and each frequency band, we simulated 20,000 time steps, and discarded the first 1000 time steps to minimise the effect of initial condition. From the remaining 19000 time steps, we calculated the proportion of duration of the network states that belongs to each energy basin.

### 2.8 Classification of individual patients based on energy values

To investigate the predictive power of pMEM energy measures, we used a support vector machine (SVM) classifier with a radial basis function (RBF) kernel and a leave-one-out cross-validation procedure to classify individual JME patients and controls. The trade-off between errors of the SVM on training data and margin maximization was set to 1. For each restingstate network and each frequency band, the feature space for classification included the energy values of all the significant local minima. In each cross-validation fold, one participant was first removed and the remaining participants’ data were used as a training set to train the classifier. To avoid over-fitting, the feature space (i.e., the local minima) was identified from the aggregated energy landscape constructed from the participants in the training set. The participant left out was then classified into one of the two groups (patients or controls). Classification performance was evaluated by the proportion of correctly classified participants over all cross-validations.

We used permutation tests to evaluate the classification results. The significance of each classification was determined by comparing the observed classification accuracy with its null distribution under the assumption of no difference between patients and controls. The null distribution was generated by 1000 random permutations of leave-one-out classification results, with group labels shuffled in each permutation. We obtained a permutation *p*-value by calculating the fraction of the permuted samples exceeding the observed classification accuracy.

## 3 Results

A summary of participant demographics and clinical characteristics is given in Table 1. The JME and control groups were well matched for age (*F*(1, 51) = 0.13, *p* = 0.72) and gender (*p* = 0.31, *χ*^2^ test). For each participant, we performed source localization of pre-processed MEG resting-state data and estimated oscillatory activity (i.e., Hilbert envelope) in each of the 90 ROIs from the AAL atlas, separately in the theta (4-8 Hz), alpha (8-12 Hz), beta (13-30 Hz), and low-gamma (35-60 Hz) bands. We focused our analysis on the differences between JME patients and controls in three resting-state networks (Fig. 2): the frontoparietal network (FPN), the default mode network (DMN), and the sensorimotor network (SMN).

**Figure 2:**
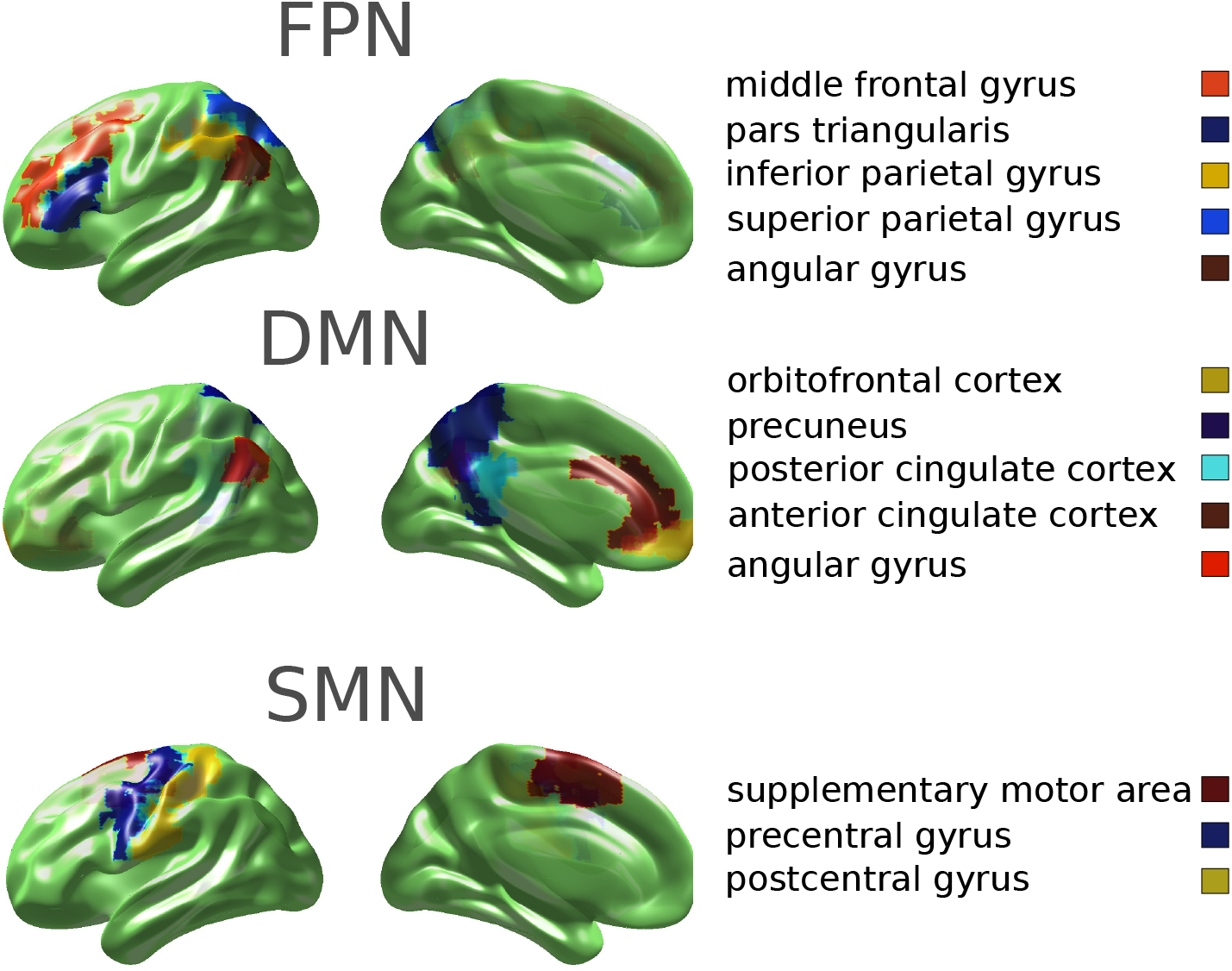
The regions of interest (ROIs) of three resting state networks: the frontoparietal network (FPN), the default mode network (DMN) and the sensori-motor network (SMN). The ROIs were obtained from the 90 AAL atlas (Hipp et al., 2012).

### 3.1 Fitting of pairwise maximum entropy models (pMEM) to MEG oscillatory activity

We thresholded an ROI’s oscillatory amplitude at each time point *t* to assign the binary states of “high” (+1) or “low” (−1) activity. The state of a network at time *t* was then represented by a binary vector, consisting of the binarised activity of all the ROIs in the network. We fitted a pMEM to the series of binarized network oscillatory activities, separately for each participant, each resting-state network, and each frequency band (Equation 3, and see Methods for details). For a network of *N* ROIs, there are a total of 2^*N*^ possible states. The pMEM provides a statistical model of the occurrence probabilities of the 2^*N*^ network states, while it satisfies the empirical constraints of mean regional activities at each ROI and pairwise co-occurrence between each pair of ROIs within the network.

To evaluate the model fit, we compared the predicted and observed occurrence probabilities of the 2^*N*^ possible network state, averaged across the participants in each group. There was a good agreement between the model predictions and observed data across networks in the JME (*R*^2^ > 0.90 in all networks and frequency bands, based on a log-log regression, Fig. 3) and control groups (*R*^2^ > 0.89). We further used an accuracy index to quantify the goodness of fit of the pMEM (Equation (7)). The accuracy index was calculated as the percentage of improvement of the pMEM fit to the empirical data compared with a null model, which assumed no pairwise co-occurrence between ROIs (i.e., an independent maximum entropy model). The pMEM achieved high accuracy indexes in both JME patients and controls (Fig. 3). A repeated-measures analysis of variance (ANOVA) on accuracy indexes showed no significant main effect of group (JME vs. controls: *F*(1, 50) = 3.40, *p* = 0.07), suggesting the robustness of the pMEM on MEG oscillatory activities in both patients and controls. There were main effects of the networks (*F*(1.44, 84.47) = 161.97, *p* < 0.000001) and the frequency band (*F*(3, 150) = 13.65, *p* < 0.000001), suggesting that the distinct properties of the networks and information carried by the frequency bands affected the goodness of fit.

**Figure 3:**
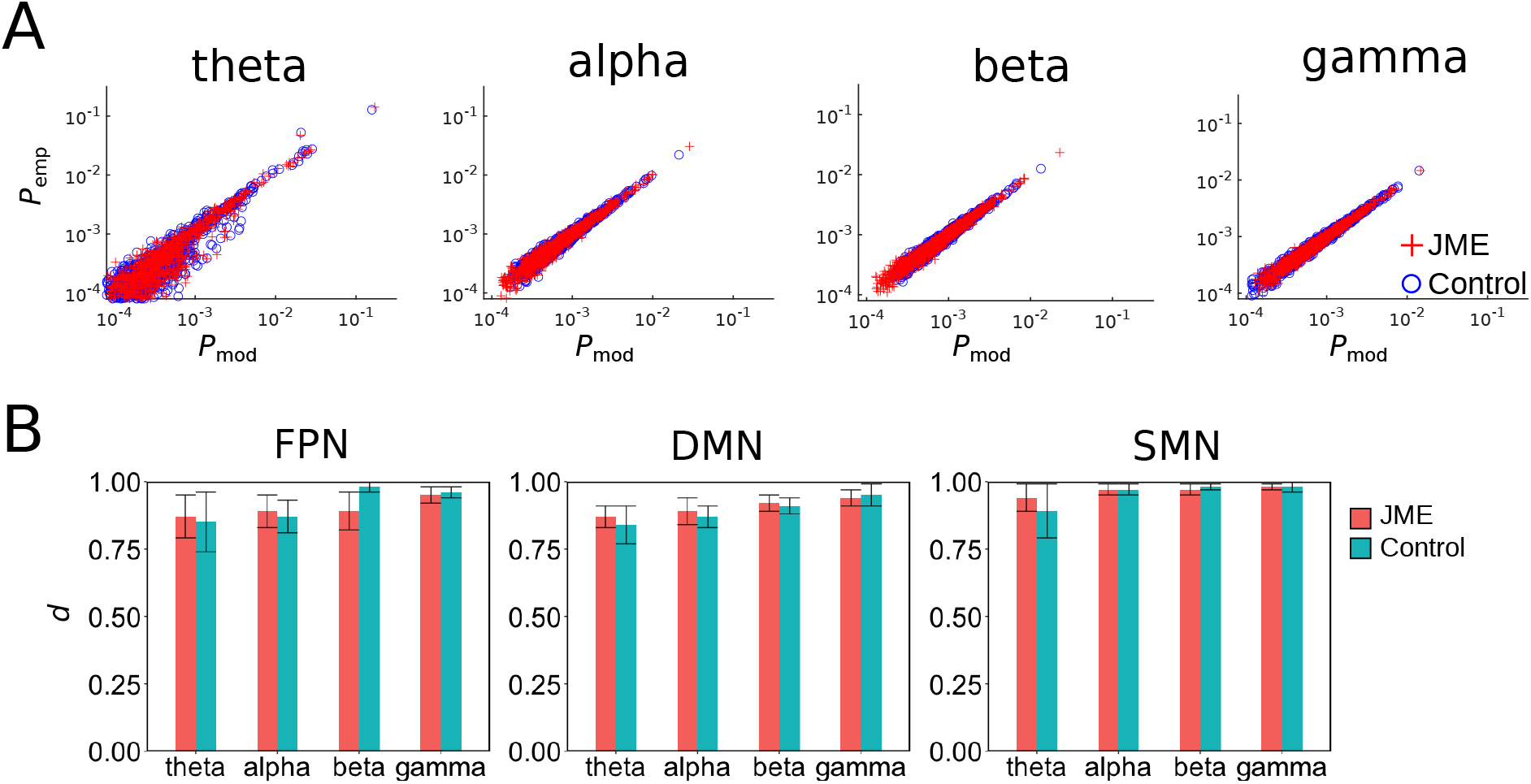
The pMEM fitting. **A**. The occurrence probability of each network state of the FPN from the fitted pMEM (*P*_mod_) was plotted against that from the empirical data (*P*_emp_). Each data point was averaged across JME patients (red) and controls (blue). **B**. The averaged accuracy index d in the JME and control groups for each network and frequency band. Error bars denote the standard errors across participants.

### 3.2 Inferences from pMEM energy landscape

The fitted pMEM yielded an energy value for each network state (Equation 4). We used energy values from the pMEM to depict an energy landscape of the network. The energy landscape is a graph representation of energy values from all possible network states (Fig. 1F). We defined two network states being adjacent if there is one and only one ROI whose binarized activity (i.e. +1 or −1) is the opposite between the two states. According to the pMEM (Equations (3) and (4)), network states with a higher energy would occur less frequently than those with a lower energy. As a result, transitions from high to low energy states would more likely to occur than that from low to high states. Here, we examined the differences in three quantitative measures of energy landscape between patients with JME and controls: (1) the number of energy minima, (2) the relative energy values at the local minima, and (3) the generative basin duration at significant minima.

#### 3.2.1 Number of energy minima

We located local minima on the energy landscape, defined as the network states with lower energy than all their adjacent states. Because a local minimum state would have a higher occurrence probability than all of its neighbouring states, transitions of network states near an energy minimum is akin to a fixed point attractor in a deterministic dynamical system, and the number of energy minima quantifies the degree of multi-stability of a network.

We calculated the number of local minima for each participant (Fig. 4) and compared it between groups, resting-state networks, and frequency bands with a repeated-measures ANOVA. Compared with controls, JME patients had significantly less local energy minima (*F*(1, 50) = 7.602, *p* = 0.008). Across all participants, there were significant main effects of the resting-state network (*F*(1.52, 76.25) = 99.89, *p* < 0.00001, Greenhouse corrected) and frequency band (*F*(2.83, 141.57) = 21.08, *p* < 0.00001). No significant network by group (*F*(1.52, 76.25) = 3.15, *p* = 0.07) or frequency band by group (*F*(2.83, 141.57) = 2.12, *p* = 0.11) interaction was observed. These results suggested that MEG oscillatory activities in JME patients had altered multi-stability in some networks and frequency bands.

**Figure 4:**
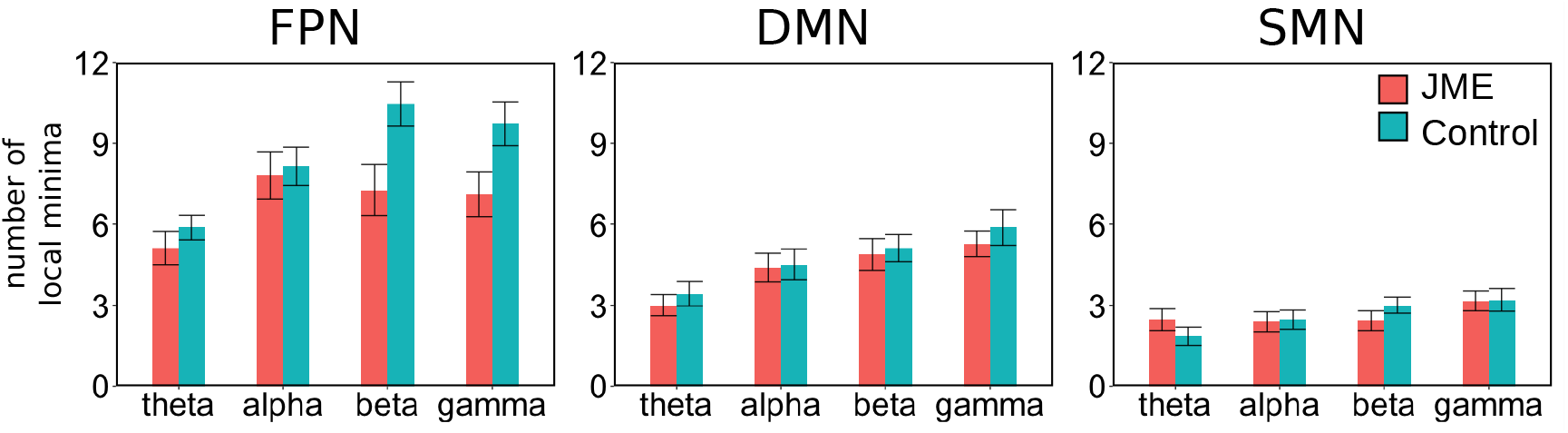
The averaged number of local minima in the JME and control groups. Error bars denote the standard errors across participants.

#### 3.2.2 Relative energy values of the local minima

To identify common energy minima at the group level, we averaged across all participants the energy value of each network state and identified the energy minima on the aggregated energy landscape. In all the three resting-state networks (FPN, DMN and SMN) and all frequency bands, permutation tests showed that the energy values of two network states, “all off” (i.e., all ROIs had low oscillatory activities [─1, −1,…, −1]) and “all on” (i.e., all ROIs had high oscillatory activities [+1, +1, …, +1]), did not differ significantly from those from a randomly shuffled energy landscape (*p* > 0.88, Bonferroni corrected). Therefore, the frequent occurrences of the “all on” and “all off” is not related to the pairwise co-occurrences between regions. In addition, the “all off” state was also the global minimum of the energy landscape, which had the lowest energy value in all network states.

For each significant local minimum state that survived the permutation test, we calculated the relative energy difference between the local minimum and the “all off” state (i.e., the global minimum) for the individual participants. Then, we compared the obtained relative energy values between the JME and control groups. This subtraction step controlled for the individual variability in the occurrence probability of the global minimum state (Watanabe et al., 2014).

In the FPN, the relative energy values at the local minima were significantly higher in JME patients than in controls in the theta-band (Fig. 5A, *F*(1, 50) = 18.90, *p* < 0.0001), beta-band (Fig. 5B, *F*(1, 50) = 15.43, *p* = 0.0002), and gamma band (Fig. 5C, *F*(1, 50) = 7.2558, *p* = 0.009), but not in the alpha band (*F*(1, 50) = 0.80, *p* = 0.37). The aggregated energy landscapes in the beta and gamma bands contained the same set of 14 local minima. Post-hoc tests showed that all the 14 local minima states had higher relative energy values in JME patients than controls in the beta band, and 5 of the 14 local minima states showed a significant group differences in the gamma band (*p* < 0.05, Šidák correction). The theta-band energy landscape contained 6 local minima states, which were a subset of the 14 local minima in the higher frequency bands, and all had higher relative energy values in JME patients than controls.

**Figure 5:**
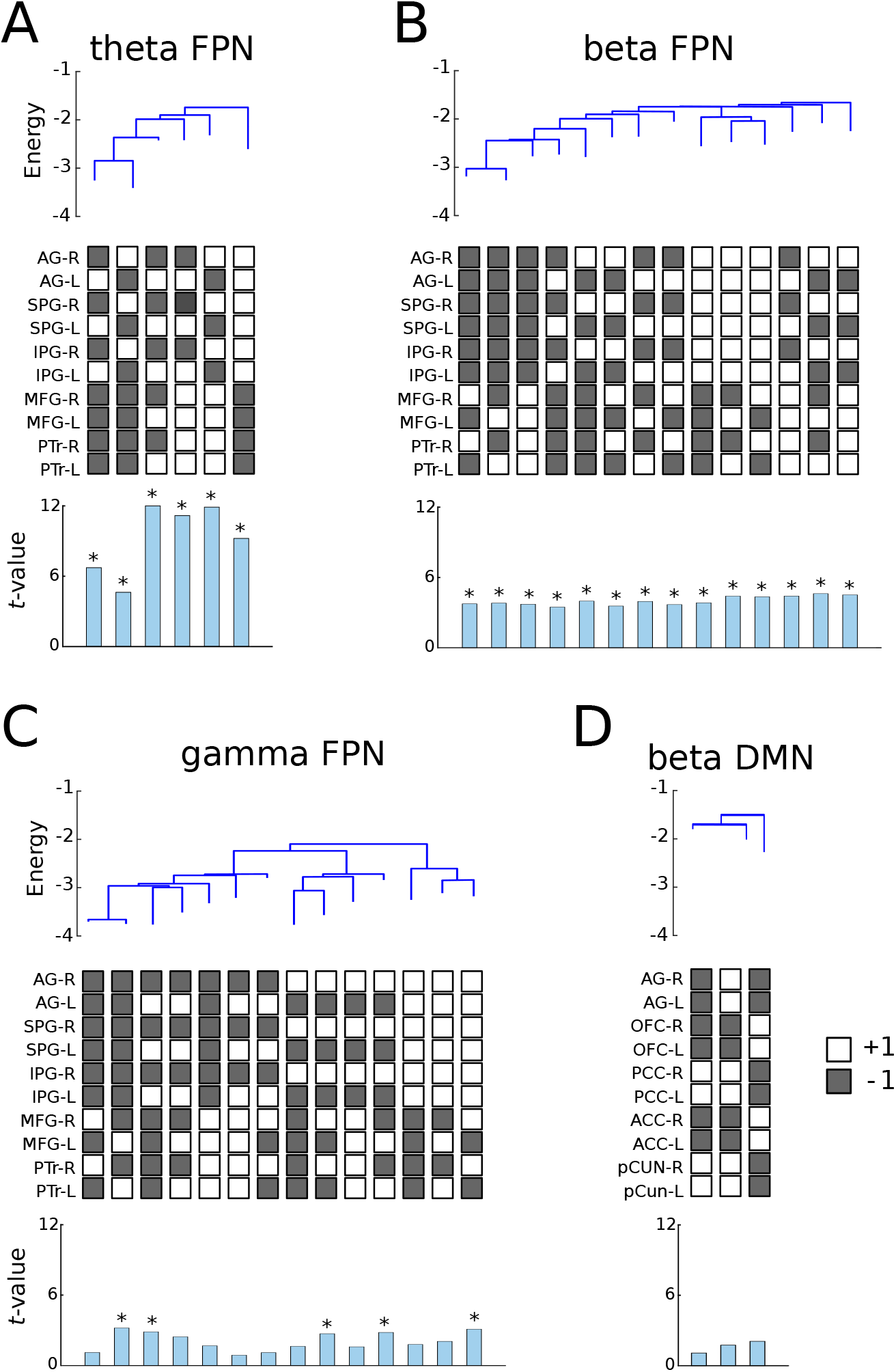
Relative energy values of the local minima in (**A**) theta FPN; (**B**) beta FPN; (**C**) gamma FPN and (**D**) beta DMN. At the top of each panel, the disconnectivity graph showed the relative energy values of local minima from the aggregated energy landscape across all participants. The end of each branch on the disconnectivity graph represent a local minimum. The middle of each panel showed the network states of the corresponding local minima. White boxes denote high oscillatory activity (i.e., a binary value of +1) and grey box denote low oscillatory activity (i.e., a binary value of −1). The bottom of each panel showed the *t*-values from two sample t-tests (JME patients vs. controls) on the relative energy values of each local minimum. Asterisks indicate significant difference between JME patient and control groups (*p* < 0.05 FDR corrected).

In the DMN, there were trends of higher relative energy values in the JME patients than controls in the beta-band (*F*(1, 50) = 3.68, *p* = 0.06) and gamma-band (*F*(1, 50) = 3.81, *p* = 0.06), and no significant difference in the theta-band (*F*(1, 50) = 0.01, *p* = 0.92) or alpha-band (*F*(1, 50) = 0.82, *p* = 0.37). One local-minima in the beta-band, comprised of co-activation in bilateral mPFC and ACC (Fig. 5D), showed a group difference in post-hoc tests at an uncorrected threshold (*t*(50) = 2.34, *p* = 0.03). In the SMN, there was no significant group difference in the relative energy values (theta: *F*(1, 50) = 1.26, *p* = 0.27; alpha: *F*(1, 50) = 0.06, *p* = 0.81; beta: *F*(1, 50) = 0.002, *p* = 0.97; gamma: *F*(1, 50) = 0.12, *p* = 0.73).

Overall, JME patients had higher relative energy values than controls in selective restingstate networks and frequency bands. This result indicates that some local minima on the aggregated energy landscape were less stable (i.e., having a higher energy level) in JME patients than controls.

#### 3.2.3 Basin duration at significant minima

Each local minimum of an energy landscape is accompanied by a basin, which includes the local minimum itself and its neighbouring states from which the local minimum is relatively easily reached (Ezaki et al., 2017). Therefore, the proportion of time for which each basin is visited gives a granular description of network dynamics. For each of the group-level significant minima on the the aggregated energy landscape, we identified all the network states belonging to the same basin. For each participant, we then used the fitted pMEM to numerically simulate network dynamics, and calculated the proportion of time for which the simulated network activities visits each basin.

In the FPN, simulations showed that network dynamics in JME patients contained shorter basin duration at those significant local minima than controls in the theta (*F*(1, 50) = 42.72, *p* < 0.000001), beta (*F*(1, 50) = 10.49, *p* = 0.002) and gamma (*F*(1, 50) = 6.18, *p* = 0.016) bands, but not in the alpha band (*F*(1, 50) = 3.92, *p* = 0.053). There was no significant group difference in the basin duration in the DMN (theta: *F*(1, 50) = 0.015, *p* = 0.90; alpha: *F*(1, 50) = 2.67, *p* = 0.11; beta: *F*(1, 50) = 2.76, *p* = 0.10; gamma: *F*(1, 50) = 3.12, *p* = 0.08) or SMN (theta: *F*(1, 50) = 0.09, *p* = 0.76; alpha: *F*(1, 50) = 0.31, *p* = 0.58; beta: *F*(1, 50) = 1.59, *p* = 0.21; gamma: *F*(1, 50) = 1.25, *p* = 0.27).

### 3.3 Classification of individual patients

We used a leave-one-out cross validation procedure for a binary classification of participant groups (JME patients and healthy controls), using the relative energy values of local minima as features. Consistent with the group comparisons (Fig. 5), the relative energy values obtained from the fitted pMEM showed significant predictive power, with high classification accuracies from theta-band FPN (92.3%, *p* < 0.001, permutation test) and gamma-band FPN (67.3%, *p* = 0.012) (Fig. 6). The classification based on the energy values from theta-band FPN achieved high specificity (89.3%) and sensitivity (94.8%). For the classification based on gamma-band FPN features, the specificity and sensitivity were 71.4% and 64.5%, respectively. The classification accuracy in the SMN, DMN and other frequency bands of the FPN was not significant (*p* > 0.26, permutation test).

**Figure 6:**
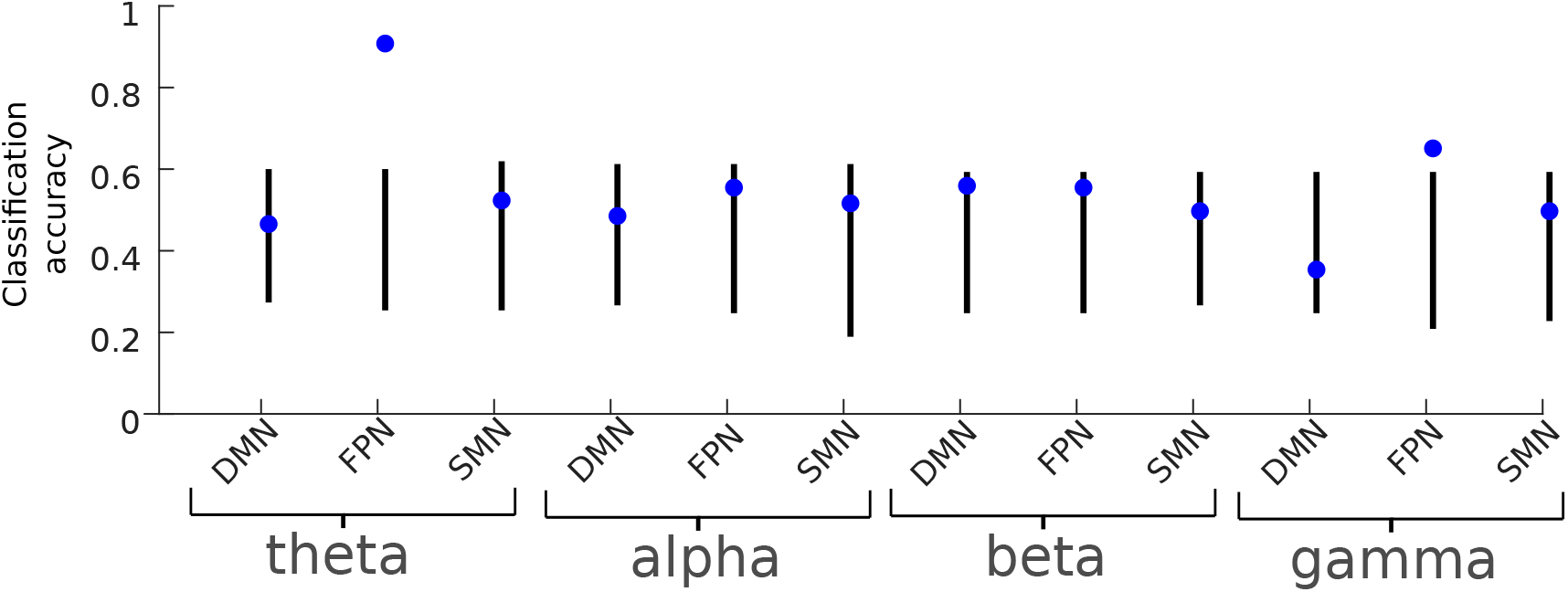
SVM leave-one-out binary classification accuracy of JME patients versus controls. The energy values of the local minima were used as features for classifiers. Blue data points denote the mean classification accuracy. Black lines denote the 95% confidence level based on 1000 permutations.

## 4 Discussion

This study applied a pMEM approach to quantify the dynamics of MEG oscillatory activity during rest. We found that patients with JME showed altered pMEM-derived energy landscapes in selective resting-state networks and frequency bands. For the energy landscapes estimated at the individual level, JME patients exhibited lower numbers of local minima than controls (Fig. 4). For the aggregated energy landscapes estimated across participants, JME elevated relative energy values at the local minima of the FPN (theta, beta, and gamma bands) oscillatory activities (Fig. 5). Our results confirmed the abnormalities of electrophysiological signals in JME (Aliberti et al., 1994), and provided new insights into JME pathophysiology affecting selective functional network configurations.

The fitted pMEM defined the energy values of all activity states of a network, from which an energy landscape of the network was depicted (Ezaki et al., 2017). Because a local minimum of the energy refers to a network state with higher occurrence probability than its neighbouring states, the fewer number of local minima and elevated energy values in JME suggested alterations in the multi-stable dynamics of the brain networks that may be prone to perturbation and ictogenesis, in line with the dynamical disease account for epilepsy (da Silva et al., 2003; Elger et al., 2000; Stam, 2005). The energy landscape further allowed to characterise clusters of energy minima and their hierarchies in terms of the disconnectivity graphs (Fig. 5). In the FPN, the energy minima with bilateral high activation in the frontal or parietal regions were clustered separately and interleaved with lateralized energy minima (i.e., high activation in unilateral ROIs). This may indicate that network states with lateralized high activation represent transition statuses between frontal and parietal dominant states. In contrast, the DMN energy minima contained co-activation in bilateral ROIs, consistent with the evidence of strong interhemispheric and long-range connectivity in the DMN during awake (Baker et al., 2014; Salvador et al., 2005).

Our results highlighted JME as a distributed network disorder involving frontal and parietal lobes (Niso et al., 2015; Fernandez et al., 2011; Wolf et al., 2015b). JME patients commonly exhibit impaired frontal cognitive functions (Piazzini et al., 2008), including working memory (Swartz et al., 1994), decision making (Zamarian et al., 2013), response inhibition (Kim et al., 2007b) and verbal fluency (O’Muircheartaigh et al., 2011). Demanding cognitive efforts during visuomotor coordination and decision-making can provoke myoclonic seizures in JME patients (Yacubian and Wolf, 2014), and the degree of cognitive dysfunctions were associated with frontoparietal BOLD fMRI activity and connectivity (Vollmar et al., 2011). Cortical and sub-cortical pathology may underlie the cognitive phenotype in JME. Activities in the lateral parietal cortex and precuneus have a dominant role in initiating and sustaining characteristic spike-and-wave discharges in JME (Lee et al., 2014). MR spectroscopy imaging of JME patients has identified reduced N-Acetyl aspartate concentrations in the frontal lobe and the thalamus (Savic et al., 2000; Zhang et al., 2016), which, together with widespread cortical morphological abnormalities (Ronan et al., 2012), indicates dysfunctions in the corticothalamic loops in JME (Hattingen et al., 2014). Further research should extend our results to associate specific abnormal energy minima to JME patients’ cognitive and behavioural phenotypes.

We further demonstrated that the pMEM and energy landscapes can be used as a generative model to simulate the duration of the network activity in each energy basins (Fig. 1) and as a predictive model for single-patient classification (Fig. 6). The normalised energies of the theta-band FPN minima achieved the best classification results (>90%), comparable with other studies (Goker et al., 2012) and consistent with our hypothesis of selective abnormalities of oscillatory activity in JME. Indeed, pathological theta oscillation were reported as a hallmark of idiopathic generalised epilepsy (Clemens, 2004), possibly owning to the involvement of the thalamus in initiating or facilitating theta oscillations through thalamocortical coherence (Sarnthein et al., 2003).

The energy landscape measures for the SMN did not significantly differ between JME patients and controls. This result might seem counter-intuitive, given that motor cortex hyperexcitability has been reported in JME (Badawy et al., 2006). Nevertheless, previous research on resting-state functional connectivity also showed the lack of altered connectivity in the motor cortex in JME (Li et al., 2015; Elshahabi et al., 2015; Liao et al., 2011). Our results suggested that the network states (i.e., patterns of co-activation) in the SMN, comprising pre- and post-central gyri as well as SMA, were not affected by JME during rest. However, this result does not rule out the possibility of network dysfunction in the motor circuit under stimulation or perturbation (Vollmar et al., 2011).

Our study provides new methods for studying the dynamics of MEG oscillatory activity. We showed that MEG oscillatory activity in resting-state networks was accurately described by the pMEM (Fig. 3) and that the model fits were comparable between JME patients and controls. The pMEM was originally developed in the field of statistical mechanics and has been applied to population of spiking neurons (Yeh et al., 2010). More recently, it has been applied to quantify the dynamics of BOLD fMRI data (Watanabe et al., 2014, 2013; Ezaki et al., 2018; Ashourvan et al., 2017; Gu et al., 2018). However, achieving satisfactory pMEM fitting requires a large number of data samples (Macke et al., 2012; Ezaki et al., 2017). Because of the low temporal resolution of the BOLD signal, the applications of the pMEM to fMRI signals often need long scanning time that may be unrealistic for clinical populations, or to concatenate data across participants that limits the possibility of individual-level inferences (Watanabe et al., 2014; Ashourvan et al., 2017). Here, we demonstrated that the high sampling rate afforded by source-localised MEG data suited well for the pMEM analysis, providing anatomically-specific and frequency-dependent results. Applying the present approach to investigate rapid changes of network energy landscapes during active tasks warrants future work.

Other methods are available to describe transient network dynamics. Microstate analysis from scalp EEG has identified successive short time periods during which the configuration of the scalp potential field remains semi-stable (Baker et al., 2014), and the spatial patterns of EEG microstates have been mapped onto distinct mental states (Brodbeck et al., 2012). Recent studies using hidden-Markov models (HMM) characterized whole-brain spontaneous activity and identified hidden states with spatiotemporal patterns at durations of 100-200 ms (Michel and Koenig, 2018; Quinn et al., 2018). Both microstate and HMM analyses are based on time-windowed approach and provide abstractions of the interactions within large-scale networks. In the current study, we defined the state of a network as an instantaneous snapshot of regional activities, and the pMEM provided a probabilistic model of the network states with minimum assumptions.

There are several limitations of this study. First, to quantify network dynamics as the occurrence probability of a finite number of network states, the oscillatory power in each ROI needed to be binarized (i.e., high vs. low activity), similar to other functional connectivity studies (Liao et al., 2010). Nevertheless, the pMEM with binarised signals showed superior performance in predicting brain structural connectivity than correlational-based methods (Watanabe et al., 2014), supporting the validity of our approach. Future research could consider more complex quantification scheme such as ternary quantization that reduces oscillatory power to ternary values (Zhu et al., 2016).

Second, the model fitting procedure for pMEM is computationally intensive. Currently, it is practically possible to fit the pMEM to a network of approximately 15 ROIs, because the number of network states increases exponentially with more ROIs. As a result, the current study focused on the dynamics within well-established large-scale resting-state networks, rather than a whole-brain network comprising all the regions. Other approximate model fitting procedures may allow us to extend our approach to larger networks with more ROIs (Ezaki et al., 2017), which is beyond the scope of the current study. To facilitate future research, we have made our analysis scripts open source and freely available^1^.

Third, the sample size in the current study is sufficient for comparing and classifying between JME patient and control groups. However, JME often exhibits as a disease with a phenotypical spectrum, with variations among seizure frequencies, epileptic traits, and treatment response (Baykan and Wolf, 2017). A larger clinical cohort with comprehensive neuropsychological assessments is necessary to investigate whether our energy landscape approach is sensitive to the quantitative spectrum of JME.

In conclusion, JME patients exhibited atypical energy landscapes of MEG oscillatory activity in selective brain networks and frequency bands, with a smaller number of local minima of the energy and elevated energy levels leading to altered multi-stable network dynamics. We further demonstrated that the pMEM and energy landscape offered generative and predictive power for discriminating between JME patients and controls. These results have the potential to be exploited in future diagnostic and pharmacological studies for a mechanistic understanding of ictogenesis in JME.

## Acknowledgements

This work was supported by the MRC/EPSRC UK MEG Partnership Grant (MR/K005464/1) and ERC (716321). DK was supported by an EPSRC PhD studentship (EP/N509449/1). KH was supported by a Clinical Research Time funding award by Health Care Research Wales during data collection for this study. BR was supported by an MRC Doctoral Training Grant (MR/K501086/1).

## Footnotes

1 https://github.com/dokato/energy_landscape

